# dP75 safeguards oogenesis by preventing H3K9me2 spreading

**DOI:** 10.1101/856021

**Authors:** Kun Dou, Yanchao Liu, Yingpei Zhang, Chenhui Wang, Ying Huang, ZZ Zhao Zhang

## Abstract

Serving as a host factor for HIV integration, LEDGF/p75 has been under extensive study as a potential target for therapy. However, as a highly conserved protein, its physiological function remains to be thoroughly elucidated. Here we characterize the molecular function of dP75, the *Drosophila* homolog of p75, during oogenesis. dP75 binds to transcriptionally active chromatin with its PWWP domain. The C-terminus IBD domain-containing region of dP75 physically interacts with the histone kinase Jil-1 and stabilizes it *in vivo*. Together with Jil-1, dP75 prevents the spreading of the heterochromatin mark–H3K9me2–onto genes required for oogenesis and piRNA production. Without dP75, ectopically silencing of these genes disrupts oogenesis, activates transposons, and causes animal sterility. We propose that dP75, the homolog of an HIV host factor in *Drosophila*, partners with Jil-1 to ensure gene expression during oogenesis by preventing ectopic heterochromatin spreading.

## INTRODUCTION

P75, also known as <u>P</u>C4 and <u>S</u>FRS1 <u>I</u>nteracting <u>P</u>rotein 1 (PSIP1) or <u>L</u>ens <u>E</u>pithelium-<u>D</u>erived <u>G</u>rowth <u>F</u>actor (LEDGF), is identified as a host factor that recruits the integrase of human immunodeficiency virus-1 (HIV-1) to targeted genomic sites (Cherepanov et al., 2003; Ferris et al., 2010; Gijsbers et al., 2010). Current model suggests that both the PWWP domain and the integrase-binding domain (IBD) from p75 are essential for tethering HIV cDNA to chromatin. While the PWWP domain finds its binding regions on host chromatin (Llano et al., 2006; Shun et al., 2008; Turlure et al., 2006), the IBD from the C-terminus region recruits HIV integrase via physical interaction (Cherepanov et al., 2004; Vanegas et al., 2005). Given its importance for HIV replication, p75 has been intensively studied for drug discovery.

Besides mediating HIV integration, the conservation of p75 across species suggests that it possesses essential molecular functions under physiological condition. In fact, depletion of pP75 in mice results in perinatal mortality, and the few ones survived into adulthood show various phenotypic abnormalities, including skeletal defects and decreased fertility (Sutherland et al., 2006). Although p75 appears to be an important factor for animal development, its molecular function is still largely unknown.

Jil-1 is an H3S10 kinase, which belongs to the serine/threonine tandem kinase family and localizes to chromatin throughout the cell cycle (Wang et al., 2001). It is functionally and structurally conserved from *Drosophila* to mammals (Jin et al., 1999; Wang et al., 2001; Zhang et al., 2003). The N-terminus of Jil-1 harbors a nuclear localization signal, followed by the tandem kinase domains in the middle. The acidic and basic domains at the C-terminus are responsible for landing Jil-1 to targeted chromatin regions (Li et al., 2013). Disrupting the function of Jil-1 leads to epigenetic changes of its targeting loci, and results in severe developmental defects, including reduced viability, segment specification defect, and maternal effect of sterility in fruit flies (Zhang et al., 2003). Despite its importance for animal development, the regulators and partners of Jil-1, which may function together with it to maintain the chromatin status, are still to be identified.

Here we identify the *Drosophila* homolog of p75–CG7946, which we refer to as *Drosophila* p75 (dP75). We show that dP75 functions as an important factor to stabilize Jil-1 and ensure fruitful oogenesis. dP75 uses its PWWP domain to associate with chromatin, and its C-terminus IBD-containing region to directly interact with and stabilize Jil-1. The dP75-Jil-1 complex protects its targeting loci from deposition of the H3K9me2, an epigenetic modification that typically leads to gene silencing (Cai et al., 2014; Deng et al., 2007; Yasuhara and Wakimoto, 2008). Accordingly, loss of either dP75 or Jil-1 leads to transcriptional suppression of their protected genes, whose expression is essential for safeguarding oogenesis and transposon silencing. Our findings provide mechanistic insights into the physiological function of the LEDGF/p75 homolog during oogenesis.

## RESULTS

### *CG7946*, the *Drosophila* homolog of p75, is essential for female fertility

To identify the *Drosophila* homolog of human p75, we performed a BLAST search of the Drosophila genome using the human p75 protein sequence. This identified *CG7946* as a single candidate, which we named as *dP75* (*Drosophila* p75). Sequence alignment revealed strong homology for both the PWWP and IBD domains (Figure 1A). In order to investigate its function, we generated two mutant alleles of *dP75* using CRISPR/Cas9 system with two different sgRNAs (Figure S1A). The *dP75^sg1^* allele lacks the ATG start codon; and the *dP75^sg2^* allele has a reading-frame shift from the 66th amino acid, resulting in a premature stop codon at the 91st amino acid (Figure S1A). To validate that both alleles are strong mutants, we generated a polyclonal antibody against the full-length dP75 protein. While we detected a strong band corresponding to the predicted size of dP75 from the control flies based on western blot, the signal was absent from *dP75 ^sg1/sg2^* flies, which are viable (Figure 1B). Immunostaining of dP75 on fly ovaries showed nucleus-localized signals in both germ cells and somatic follicle cells. These signals were undetectable in *dP75^sg1/sg2^* mutants (Figure 1B). These data suggest that neither mutant could produce functional proteins, and thus likely serve as null alleles. To avoid any effects from the potential secondary mutations resulting from CRISPR off-targeting, we used trans-heterozygotes, *dP75 ^sg1/sg2^*, in our study and hereafter refer to them as *dP75* mutants.

**Figure 1.**
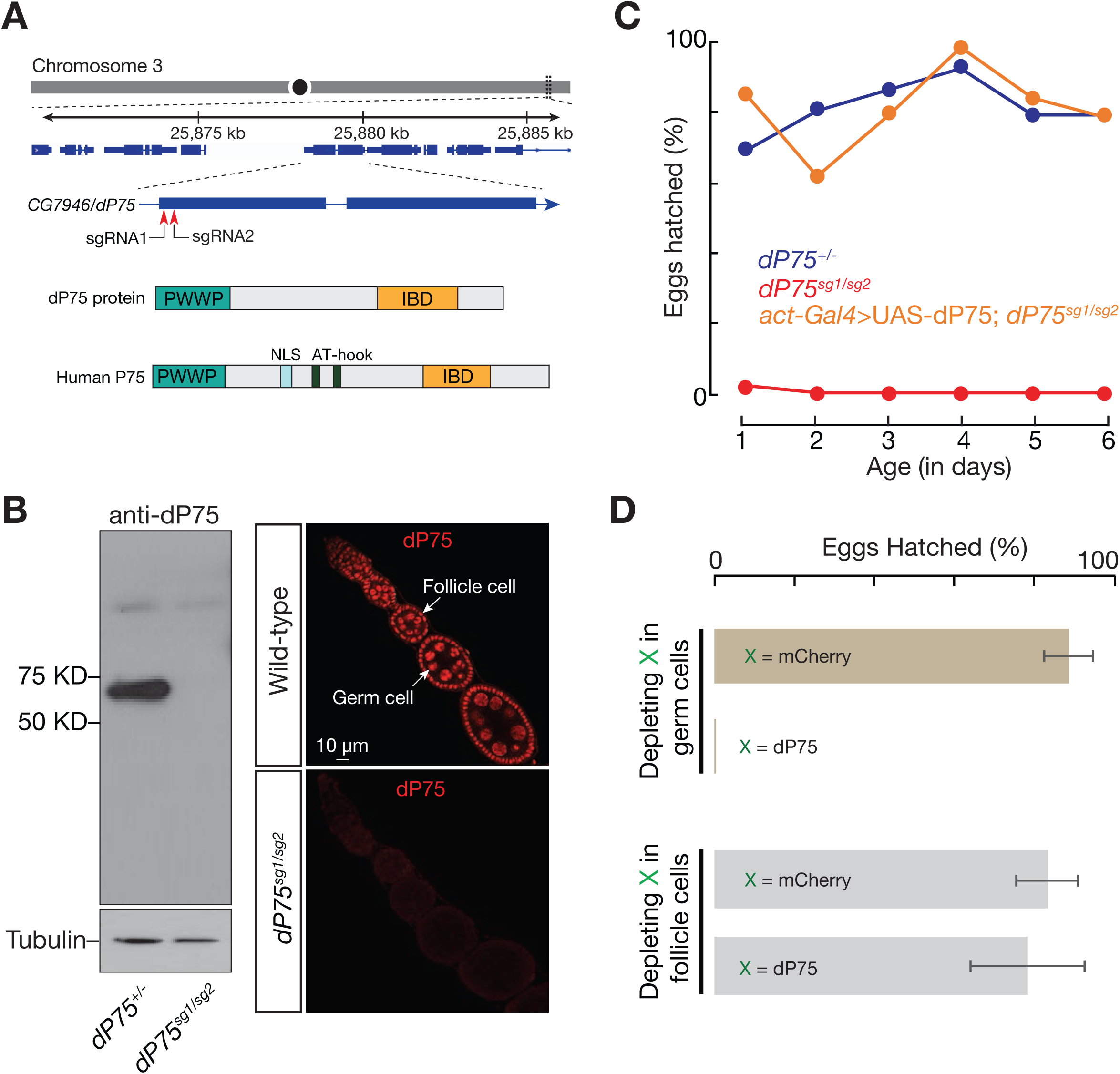
CG7946/dP75, the LEDGF/p75 homolog in Drosophila, is essential for female fertility. (A) dP75 resides on the right arm of chromosome 3, encoding a protein with PWWP and LEGDF domains. IBD = Integrase-binding domain. Two sgRNAs used to generate dP75 mutants are labelled. (B) Characterization of dP75 mutants. Left panel: western blot of ovarian lysate using polyclonal antibody against dP75. The bright band between 50kD and 75kD corresponds to the predicted size of dP75 protein. Right panel: immunostaining with dP75 antibody in wild-type and dP75 mutant ovaries. (C) Fertility of females over time. Reintroducing dP75 transgene driven by actin-Gal4 into dP75 mutant rescued the sterility phenotype. (D) Fertility of females with dP75 depletion in either germ cells or in somatic follicle cells. shRNA against mCherry was used as a control.

Loss of dP75 has no effect on fly viability, but led to female sterility (Figure 1C). Additionally, reintroducing dP75 into mutants through a transgene fully rescued the fertility defects (Figure 1C), arguing that dP75 serves as a bona fide factor to maintain female fertility. Next, we sought to determine whether the sterility phenotype of *dP75* is caused by its loss of function in germ cells or somatic follicle cells. To achieve this, we employed cell-type specific RNAi assays (Figures S1B-S1E). While only depleting dP75 in follicle cells had a marginal effect on fertility, specifically suppressing dP75 in germ cells by RNAi recapitulated the sterility phenotype from the mutants (Figures 1D and S1E). Our data thus suggest that dP75 is indispensable in germ cells to maintain fruitful oogenesis.

### dP75 binds to actively transcribed genomic regions in fly ovaries

Since the PWWP domain is often involved in chromatin binding (Qin and Min, 2014), we next tested whether dP75 binds to any specific genomic regions in fly ovaries. To facilitate this study, we generated a fly allele that harbors one V5 tag sequence at the endogenous *dP75* locus, thus producing V5::dP75 proteins (Figures 2A and 2B). Adding V5 tag appears to have negative effect on dP75 protein function and localization (Figure 2A). We next performed chromatin immunoprecipitation followed by high-throughput sequencing (ChIP-Seq). Our ChIP-seq data showed that dP75 preferentially covers the genes engaged in active transcription (Figures 2C and 2D). In fly ovaries, there were 6985 genes producing transcripts with the FPKM (Fragments Per Kilobase of transcript per Million mapped reads) ≥ 3. Among them, 6575 (94.1%) showed significant enrichment for dP75 binding (Figure S2A). On the contrary, for the 9706 genes having an FPKM value lower than 3 in fly ovaries, only 2092 (21.5%) of them were enriched by dP75 binding (Figure S2A). Additionally, we calculated the enrichment of dP75 peaks over intergenic regions, 5’ and 3’UTRs, and exonic and intronic regions respectively. While intergenic regions had minimum, if any, binding of dP75, it showed strong enrichment over exons and the 5’ and 3’ UTRs (Figure S2B).

**Figure 2.**
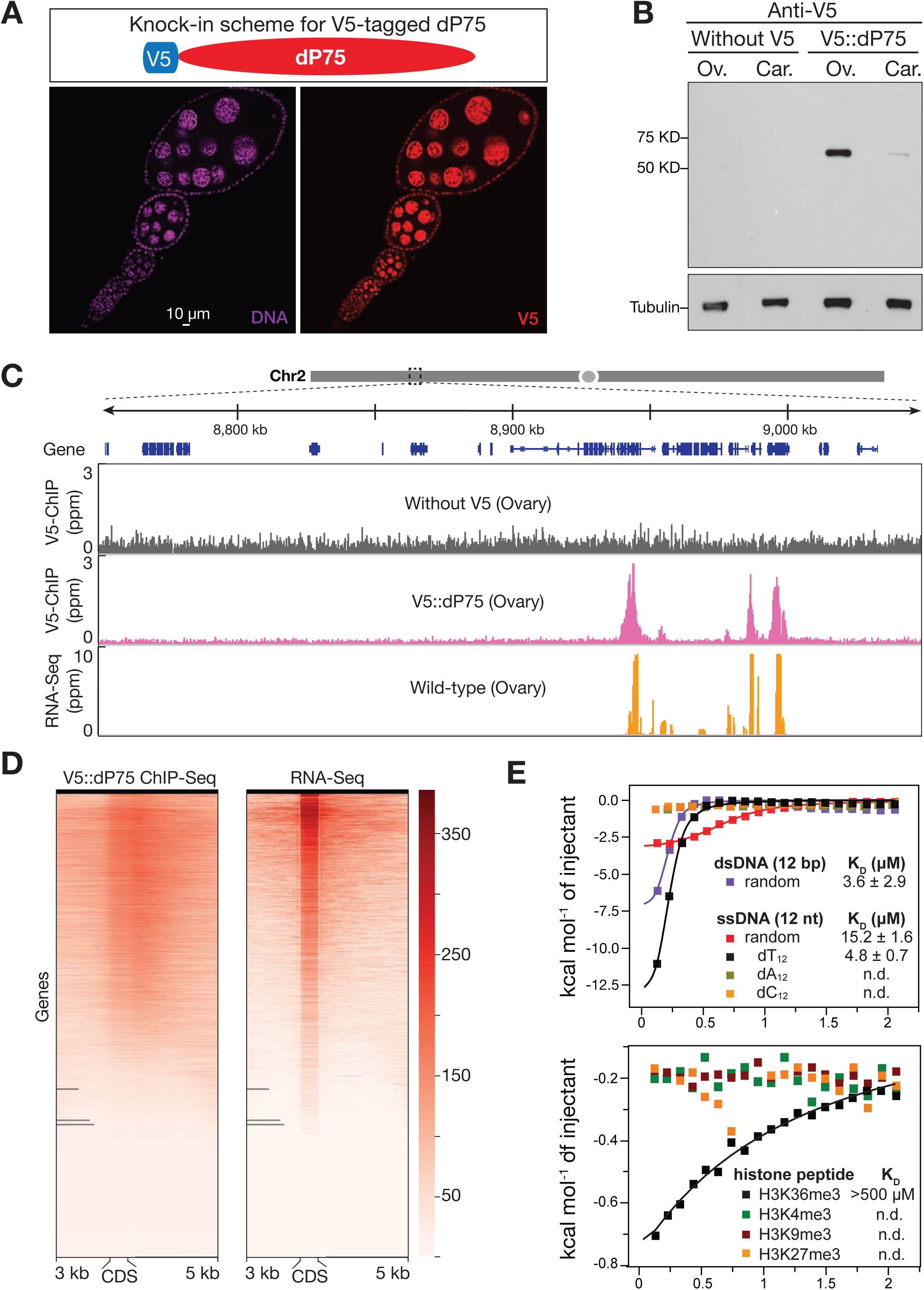
dP75 binds to actively transcribed chromatin regions in vivo. (A) dP75 expression pattern as examined by V5 antibody staining in ovaries with V5 knocked-in at endogenous locus of dP75. (B) Validation of V5 tag knock-in at endogenous dP75 locus by western blot. Car = Carcass: fly body with ovaries ectomised. (C) V5 ChIP-Seq and RNA-Seq signal across a 300 kb representative genome region on chromosome 2. V5 ChIP-seq performed in ovaries without V5 tag serves as negative control. (D) Heatmap for V5-dP75 ChIP-seq signal on genes and their mRNA levels in wild-type fly ovaries. Each line from the heatmap represents the coding region of one gene plus 3kb upstream and 5kb downstream of the coding region. (E) Isothermal titration calorimetry (ITC) measurement of the binding affinity of PWWP domain to different oligos (upper panel), and various histone peptides (lower panel).

To explore how dP75 is recruited to its targeting loci on chromatin, we employed Isothermal titration calorimetry (ITC) to measure the binding affinity of purified PWWP domain to a few potential substrates: single-stranded DNA, double-stranded DNA, peptides of H3K4me3, H3K9me3, H3K27me3, and H3K36me3. The PWWP domain of dP75 had high binding affinity with double-stranded DNA (K_D_ = 3.6±2.9 µM) and Oligo-dT single-stranded DNA (K_D_ = 4.8±0.7 µM) (Figure 2E). When using different modified histone peptides as substrates, it did not exhibit any binding preference with them except for a low affinity with H3K36me3 (K_D_ > 500 µM, Figure 2E), a histone marker for transcriptionally active genes (Kolasinska-Zwierz et al., 2009). Our in vitro data suggest that dP75 is recruited to chromatin by H3K36me3 and this binding might be further enhanced by interacting with DNA.

### dP75 physically interacts with histone kinase Jil-1 both *in vivo* and *in vitro*

Given that the IBD (Integrase Binding Domain) of mammalian p75 often mediates protein interactions (Tesina et al., 2015), we performed V5 IP-mass spectrometry to search for the binding partners of dP75 in V5::dP75 fly ovaries. From the V5::dP75 ovary lysate, we recovered 149 peptides from dP75, which was listed as the top hit from mass spectrometry data. In control lysates derived from the flies without a V5 tag, we did not detect any peptides derived from dP75, highlighting the high stringency of our IP condition (Figure 3A). Besides dP75, the second strongest hit was from histone kinase Jil-1 in V5::dP75 lysate (with 14 peptides being detected), but none from the control (Figure 3A). Supporting our findings, a previous study in *Drosophila* S2 cells also identified Jil-1 and CG7946/dP75 in the same complex (Wang et al., 2013). Accordingly, we also noticed that Jil-1 and dP75 colocalize in the nucleus of both germ cells and somatic follicle cells in fly ovaries (Figure 3B).

**Figure 3.**
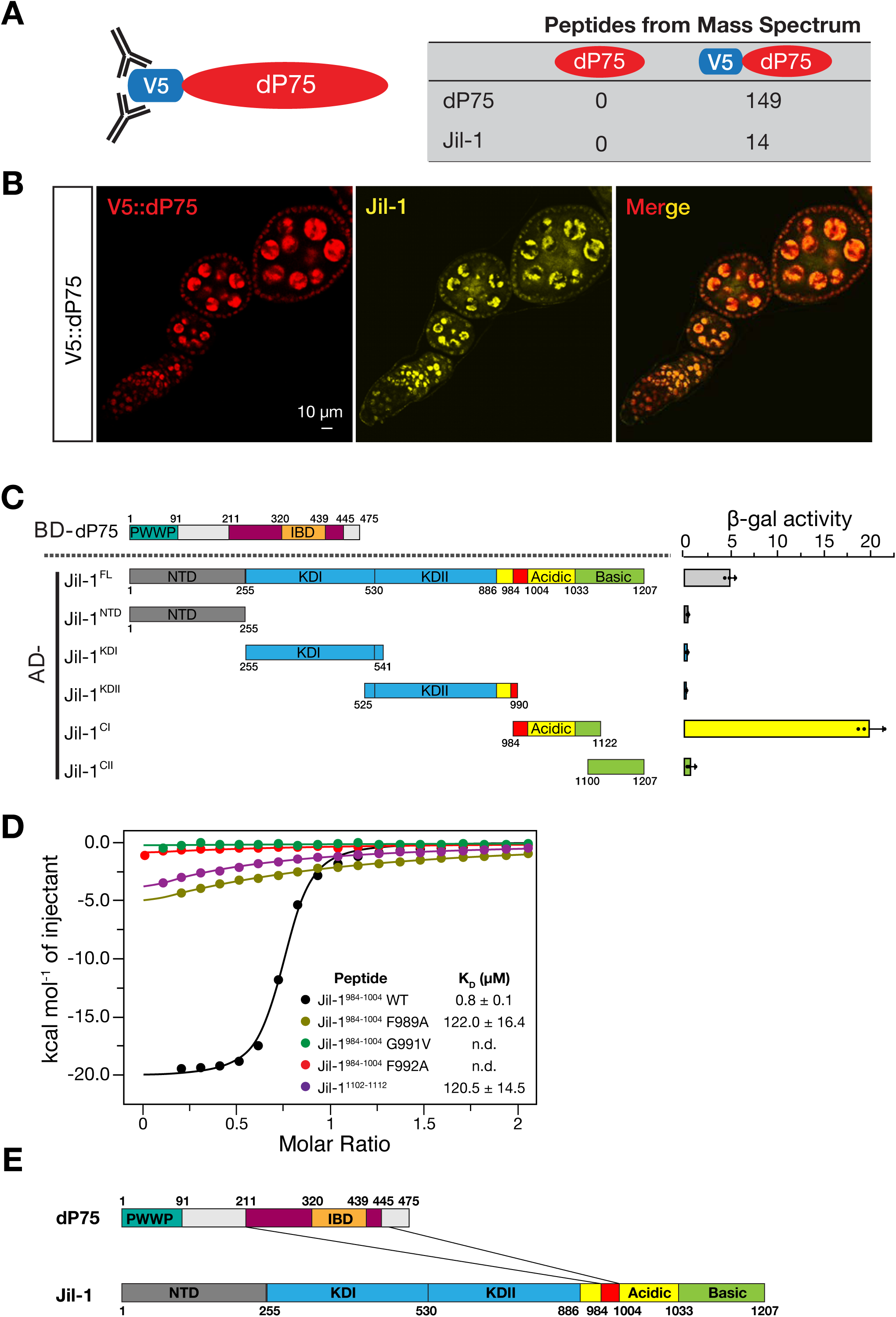
dP75 interacts with Jil-1 both in vivo and in vitro. (A) Identification of dP75 interacting proteins by affinity purification and mass spectrometric analysis. The number of peptides identified from mass spectrometry for dP75 and Jil-1 in each sample is listed. (B) dP75 and Jil-1 colocalize in fly ovaries. (C) Interaction measurement between dP75 and full-length or truncated versions of Jil-1 by yeast-two-hybrid assay. Interaction strengths are indicated by -gal activity. Jil-1C1 fragment exhibits the strongest interaction with full length dP75. (D) Isothermal titration calorimetry (ITC) to measure the interaction between dP75 and wild-type or mutated Jil-1 peptide (984-1004 fragment). Mutation of the conserved amino acids in the FYGF region of Jil-1 peptide severely attenuates its binding affinity with dP75. (E) Graphic illustration for the interacting regions between dP75 and Jil-1.

We next tested whether the interaction between dP75 and Jil-1 is direct. To do this, we used a yeast two-hybrid assay (Y2H) with the beta-galactosidase activity as a readout. Full-length dP75 showed interaction with either the full-length Jil-1 or the C-terminal fragment of Jil-1 (residues 984-1122), but not the N-terminal domain or the kinase domain (Figure 3C). We then sub-divided dP75 and tested the interaction between each fragment and the C-terminus region of Jil-1 to further map the interaction region on dP75 by Y2H. The C-terminus fragment including the IBD of dP75 was shown to be responsible for binding with Jil-1 (Figure S3).

Human P75 has been reported to interact with its cellular binding partners via a conserved (E/D)-X-E-X-F-X-G-F motif on these proteins (Tesina et al., 2015). Two regions containing the FYGF sequence are found within the C-terminal fragment of Jil-1, which are located at the proximal site (residue 989-992) and the distal site (residues 1107-1110), respectively. To validate the interactions between dP75 and Jil-1, we performed isothermal titration calorimetry (ITC) using different Jil-1 peptides. The binding to the peptide from the proximal site (EVQADFYGFDE; residues 984-1004) is 150-fold stronger (K_D_ = 0.8 ± 0.1 μM) than to the peptide at the distal site (KREENFYGFSK; residues 1102-1112; K_D_ = 120.5 ± 14.5 μM). Alanine substitution of the key residues (F989A, G991V, and F992A) dramatically reduced the binding (Figure 3D). Taken together, our data indicate that in fly ovaries, dP75 uses its IBD-containing C-terminus region to recruit the histone kinase Jil-1 by directly interacting with its C-terminus (Figure 3E).

### dP75 is required to stabilize Jil-1 protein *in vivo*

Since dP75 possesses chromatin binding property and physically interacts with Jil-1, we next tested whether Jil-1 localization is dP75 dependent. Either depleting or mutating dP75 resulted in a dramatic reduction of the endogenous Jil-1 in the nucleus (Figures 4A and S4A). To further validate this dependency, we next generated mosaic ovaries containing *dP75* mutant cells in adjacent to dP75 heterozygotes or wild-type cells within the same ovariole (Figure 4B). This ensures all cells were treated and imaged under identical conditions. As shown in Figure 4C, Jil-1 staining gave rise to nuclear foci in the cells producing dP75 (marked by the presence of GFP). In contrast, *dP75* mutant cells always displayed very weak, if any, Jil-1 signals (Figure 4C). To validate this result, we introduced Flag::Myc tagged Jil-1 transgene under a germline specific driver to either wild-type or ovaries with dP75 depleted in germ cells, and performed both immunostaining and western blot to detect the expression of transgenic Jil-1 protein. Results from both experiments showed that without dP75, the levels of transgenic Jil-1 were reduced to nearly undetectable (Figures 4D and 4E). Because loss of dP75 (either by RNAi depletion or under chromosomal mutation) had no influence on the steady state level of Jil-1 mRNA (Figures 4F and S4B), these results indicate that dP75 is required to stabilize Jil-1 protein in vivo.

**Figure 4.**
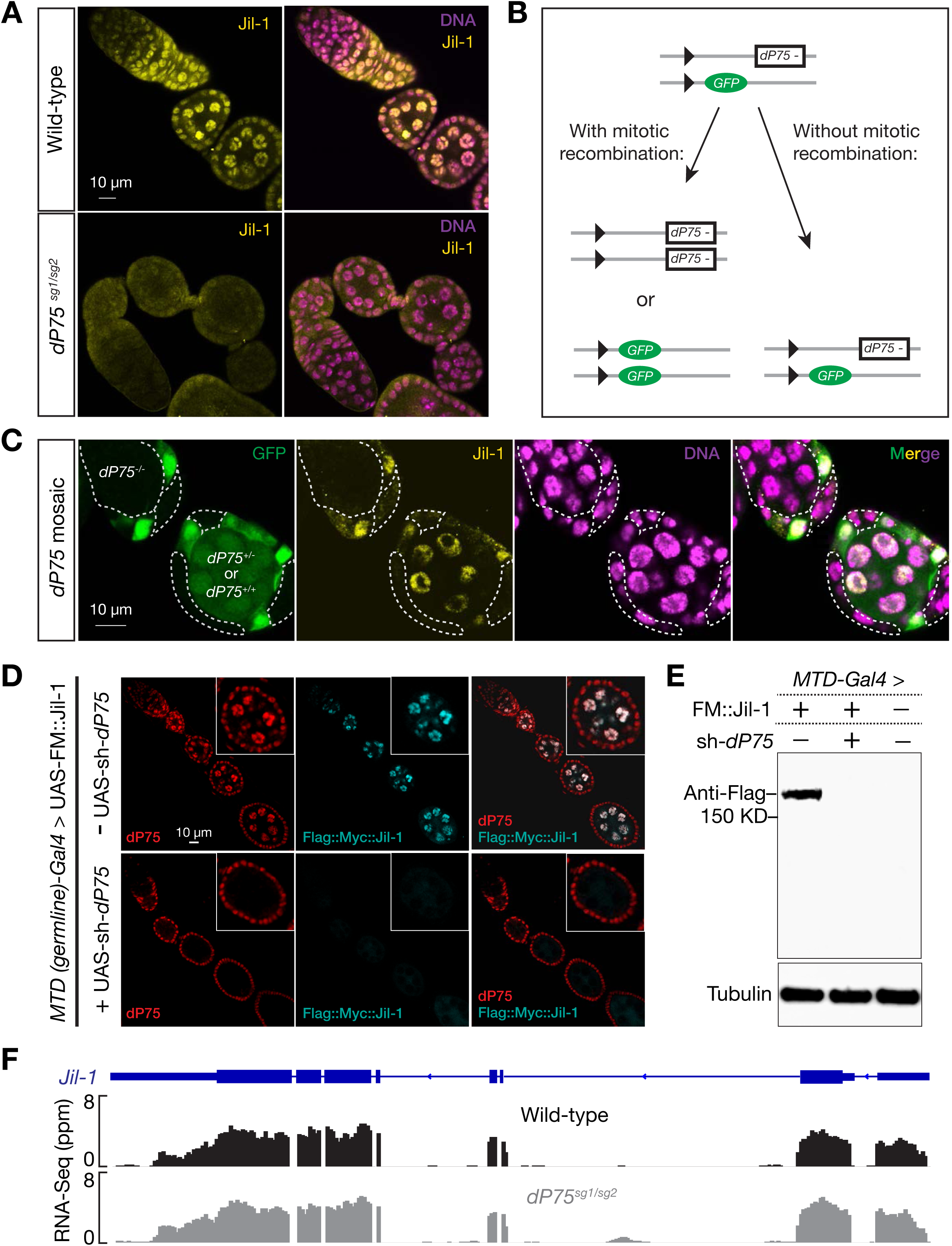
dP75 is required for stabilizing Jil-1 protein. (A) Immunostaining of Jil-1 in the ovary from either wild-type or dP75 mutants. Jil-1 shows robust nucle-us-localized signal in wild-type flies, while these signals are nearly absent in dP75 mutant. (B) Strategy for mosaic analysis in (C). (C) clonal analysis of Jil-1 expression. dP75 mutant clones were generated through strategy shown in (B) and were subjected to immunostaining with Jil-1 antibody seven days after clone induction. dP75 mutant clones are marked by absence of GFP, and outlined with dashed lines. (D) Immunostaining of transgenic Jil-1 protein in either control or in ovaries with dP75 depleted in germ cells. UAS-Flag::Myc::Jil-1 driven by germline specific Gal4 shows robust expression in wild type ovaries, but is undetectable when dP75 is depleted. (E) Western blot for transgenic Jil-1 protein in control and dP75 depleted ovaries. (F) RNA-Seq data show that the abundance of jil-1 mRNAs is not affected in dP75 mutants.

### dP75 restrains H3K9me2 spreading

Previous studies suggest that Jil-1 antagonizes the H3K9me2 in salivary glands (Deng et al., 2007; Zhang et al., 2006). We next investigated whether dP75 is responsible for restraining H3K9me2 during oogenesis. In wild-type fly ovaries, immunostaining of H3K9me2 showed limited bright puncta in the nucleus (Figure 5A). Loss of dP75 appears to lead to broader deposition of H3K9me2 in ovarian cells (Figure 5A), suggesting the spreading of H3K9me2. To test whether dP75 indeed inhibits H3K9me2 spreading with base-pair resolution, we performed ChIP-seq experiments. In wild-type fly ovaries, we detected strong enrichment of H3K9me2 modification at the pericentromeric regions, consistent with previous findings (Yasuhara and Wakimoto, 2008). However, in *dP75* mutant ovaries, the intensity of H3K9me2 peaks decreased drastically at the pericentromeric regions, while we detected more even deposition across chromosome arms (Figure 5B).

**Figure 5.**
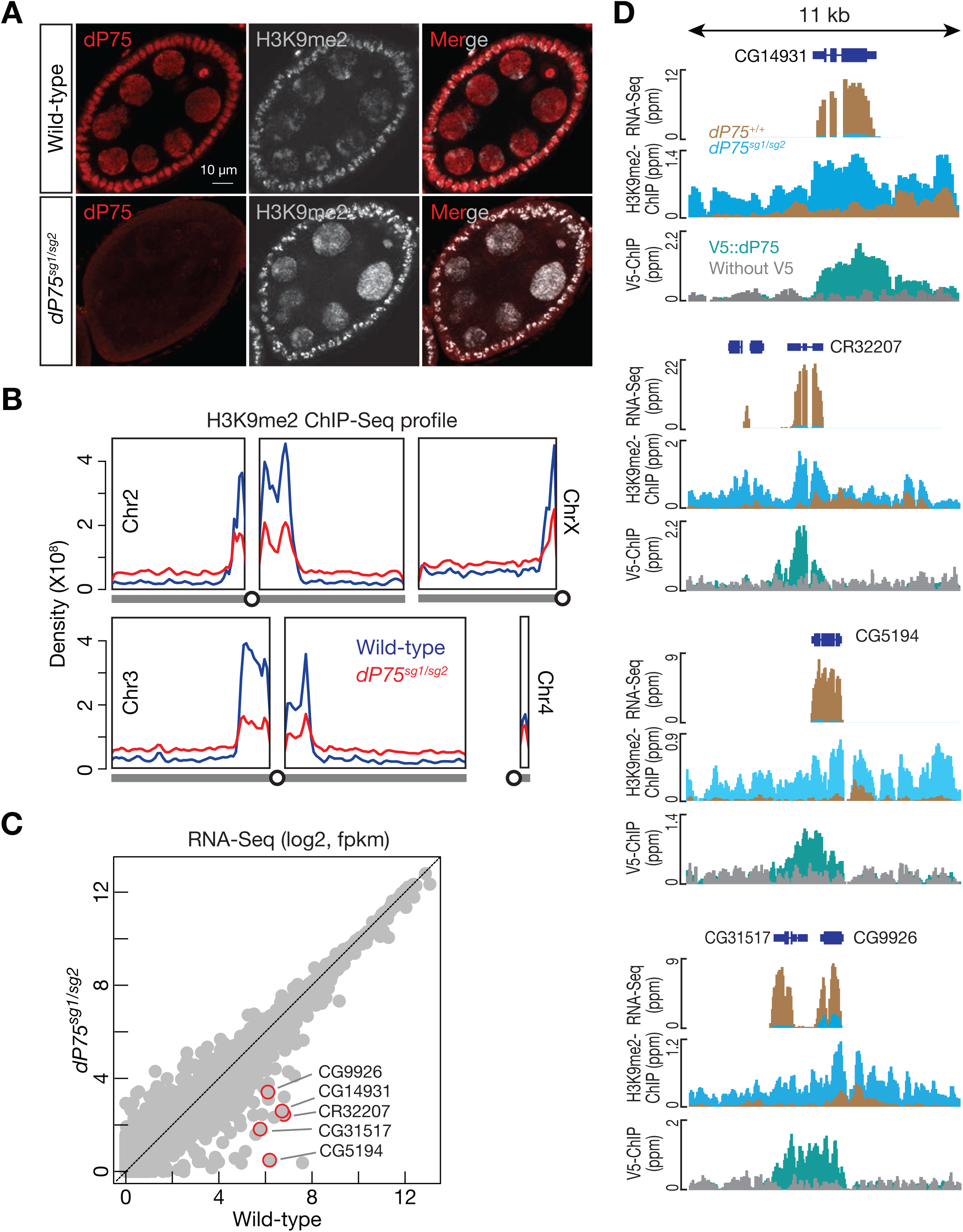
dP75 restrains H3K9me2 from spreading on chromosome arms. (A) Co-immunostaining of dP75 and H3K9me2 in wild-type or dP75 mutant ovaries. (B) Density of H3K9me2 ChIP-Seq signal across chromosome arms in wild-type or dP75 mutant ovaries. The same sequencing depth was used for analysis of the two samples. (C) Expression of genes measured by RNA-Seq. (D) Browser view of RNA-Seq, H3K9me2 occupancy, and dP75 ChIP-seq signals for the top decreased genes in dP75 mutants. Loss of dP75 leads to increase of H3K9m32 deposition and gene silencing.

Given that H3K9me2 deposition often promotes gene silencing (Caro et al., 2012), we next performed mRNA sequencing (RNA-Seq) to determine whether loss of *dP75* affects gene expression. In *dP75* mutant ovaries, we detected 74 genes that decreased their mRNA abundance by more than 3-fold (Figure 5C). Among them, 37 also showed > 3-fold reduction upon depleting dP75 only in germ cells (Figure 6A). Closer examination of the top decreased genes on the genome-browser revealed that in wild-type fly ovaries, dP75 binding prevents the deposition of H3K9me2 onto them (Figure 5D). Upon loss of dP75, they were coated by H3K9me2, and ectopically suppressed at the transcriptional level (Figure 5D).

**Figure 6.**
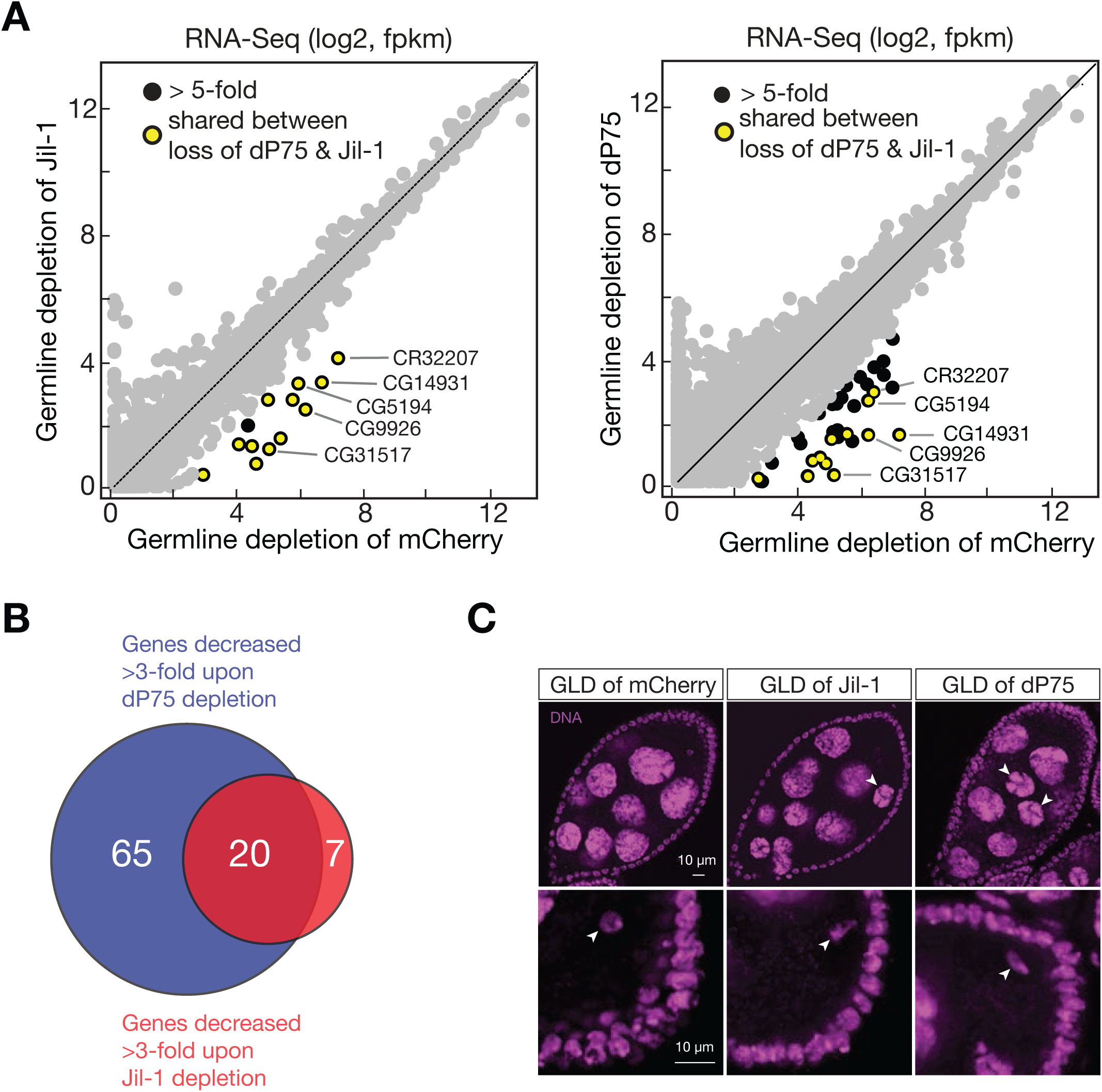
dP75 and Jil-1 function together to ensure oogenesis. (A) Expression of genes measured by RNA-Seq. (B) dP75 and Jil-1 protect common targets for transcription. (C) DAPI staining shows the similar developmental defects of germ cells by either depleting of dP75 or Jil-1 in germ cells. Upper panel: arrow heads point the nurse cells failed to pass the five-blob configuration. Lower panel: oocyte DNA failed to compact into karyosome (pointed by arrow head).

### dP75 and Jil-1 function together to ensure oogenesis

Given the interactions between dP75 and Jil-1, we next asked whether Jil-1 has similar impacts on oogenesis. Depleting Jil-1 in germline cells also led to the spreading of H3K9me2 on chromosomes (Figures S5A-S5C). Additionally, our transcriptome analysis revealed that the same group of genes was suppressed upon loss of Jil-1 or dP75 (Figures 6A and 6B). Without Jil-1 being produced in ovarian germ cells, we detected 13 genes significantly decreased for > 5-fold at mRNA level (Figure 6A). Among them, 12 were also suppressed when dP75 was absent (Figure 5C and 6A). Similar as dP75, Jil-1 from the germline cells is essential for the maintenance of animal fertility (Figure S5D).

Phenotypically, depleting dP75 or Jil-1 in germ cells resulted in similar oogenesis defects: irregular sized egg chambers along the ovariole; more than 15 nurse cells encapsulated in one egg chamber; five blob nurse cell chromosome structure in late stage egg chambers (Figure 6C and Table S1). Moreover, we also observed severe karyosome defects when dP75 or Jil-1 was absent from germ cells. During oogenesis, oocyte chromosomes arrested at the meiosis prophase I stage were packaged into a subnuclear structure named karyosome, appearing as a condensed round structure (Staeva-Vieira et al., 2003). In control ovaries, almost all of the stage 7-8 egg chambers had normal karyosome (98%, n = 56). However, loss of dP75 in germ cells resulted in 27% of oocytes in these stages having stretched karyosome (n = 55; Figure 6C). The defective karyosomes were also frequently observed in ovaries upon Jil-1 depletion (34% of stage 7-8 egg chambers, n = 92; Figure 6C). All the above results indicate that dP75 and Jil-1 function together on maintaining fruitful oogenesis.

### dP75 is required for transposon silencing

From our RNA-Seq data, we noticed that BoYb, a piRNA component (Handler et al., 2011), is among the top targets protected by dP75 and Jil-1 (Figure 7A, Figure S6A). Its mRNA decreased 95% and 80%, respectively, upon dP75 or Jil-1 depletion in fly ovaries (FPKM: 28.78 in control ovaries, 1.58 in dP75 depleted ovaries, and 5.74 in Jil-1 depleted ovaries). In fact, dP75/CG7946 was previously recovered in a genome-wide screen identifying piRNA pathway components (Czech et al., 2013). Given the importance of BoYb in transposon silencing in germ cells (Handler et al., 2011), we measured transposon transcripts in fly ovaries when dP75 is absent. Our RNA-Seq experiments showed that 21 transposon families increased their transcripts by more than 2-fold in *dP75* mutants (Figure 7B). The increasing of transposon mRNA was further validated by RNA FISH (Fig 7C). Consistently, transposon transcripts also increased upon loss of Jil-1 in germ cells (Figure S6B), including Burdock and HMS-Beagle, which were the most affected transposon families in *dP75* mutants. Taken together, our data indicate that the dP75-Jil-1 complex is essential for transposon suppression during oogenesis, likely by promoting the expression of key piRNA pathway components, such as BoYb (Figure 7D).

**Figure 7.**
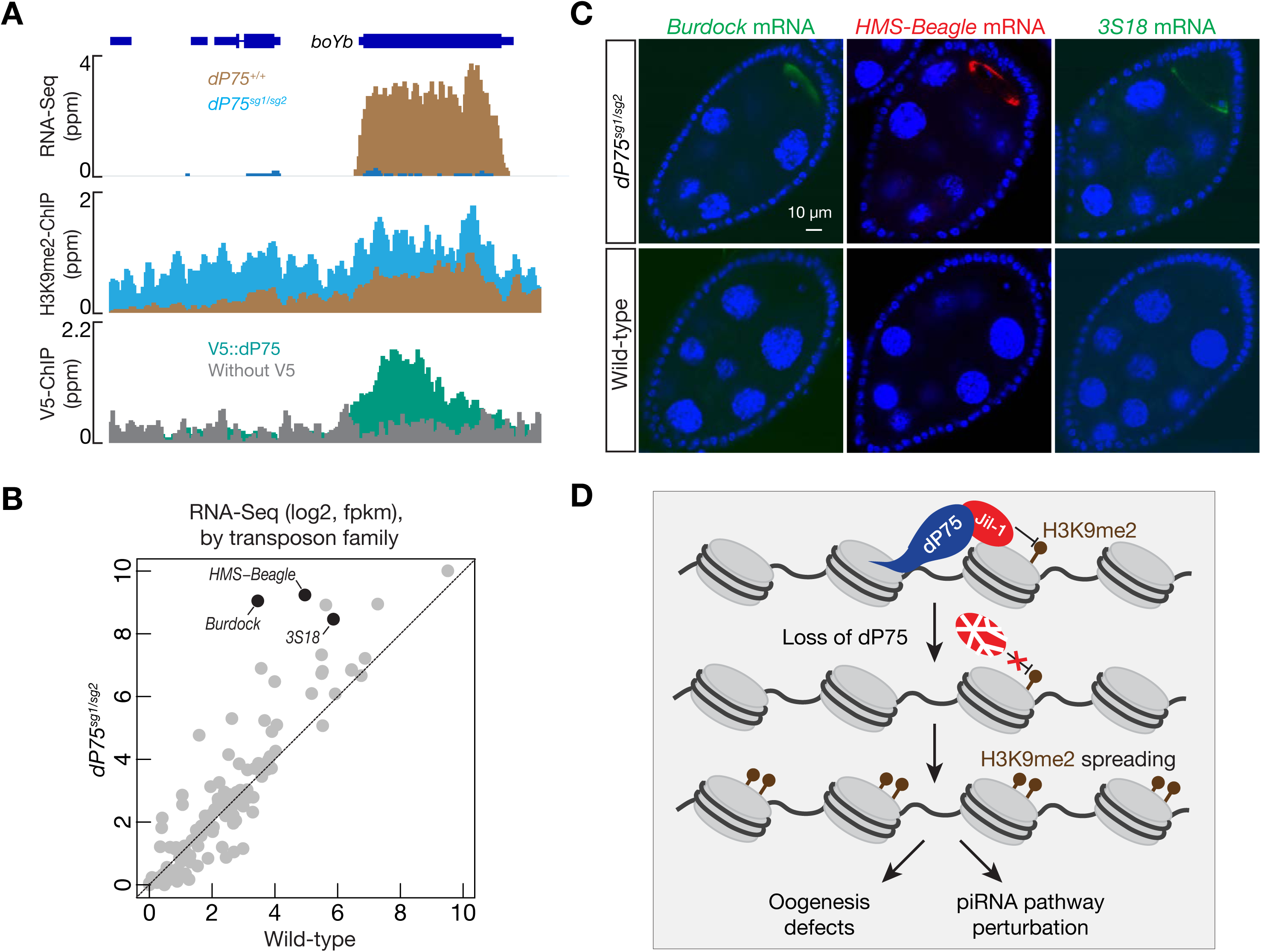
dP75 is required for transposon silencing during oogenesis. (A) RNA-Seq, V5::dP75 ChIP-Seq, and H3K9me2 ChIP-Seq signals across the *boYb* gene, which encodes a piRNA pathway component for transposon silencing. dP75 protects *boYb* from H3K9me2 spreading. (B) The change in transposon expression, relative to control, was compared for *dP75* mutants. (C) RNA FISH to detect the expression and localization of transposon transcripts. (D) Graphic model to depict the function of dP75 during oogenesis. dP75 physically interacts with Jil-1, antagonizing the spreading of H3K9me2. Upon loss of dP75, Jil-1 protein is unstable, and leads to H3K9me2 deposition on genes that are required for ensuring oogenesis and transposon silencing.

## DISCUSSION

By studying the physiological function of the homolog of mammalian p75 (dP75), here we uncover its molecular role on ensuring fruit fly oogenesis. Our study reveals that dP75 binds to actively transcribed chromatin via its PWWP domain. Through the IBD-containing C-terminus region, it recruits and stabilizes Jil-1, a histone kinase for the phosphorylation of the 10^th^ residue of Histone 3 (H3S10) (Wang et al., 2001; Zhang et al., 2006). Previous findings suggest that H3S10 phosphorylation antagonizes the methylation of H3K9, and thus counteracting gene silencing (Bao et al., 2007; Zhang et al., 2006). Consistently, dP75-Jil-1 binding appears to block the spreading of H3K9me2, thus promoting the expression the genes that are essential for oogenesis and transposon silencing (Figure 7D).

In humans, p75 interacts with chromatin regions that are actively transcribing (De Rijck et al., 2010), thus facilitating HIV integrating into these loci (Ferris et al., 2010; Vanegas et al., 2005). Human p75 harbors one PWWP domain and two AT-hooks, all of which appear to contribute to target binding (Llano et al., 2006). The PWWP domain of human p75 shows binding affinity to mononucleosomes with trimethylated Histone 3 at lysine 36 (H3K36me3) in vitro (Eidahl et al., 2013), which usually marks actively transcribed regions (Bannister et al., 2005; Barski et al., 2007). As its name indicates, the two AT-hooks can directly bind to the minor groove of adenine-thymine (AT) enriched DNA (Llano et al., 2006; Reeves and Nissen, 1990). Therefore, with the PWWP domain and AT hooks functioning cooperatively, human p75 can simultaneously interact with both modified histones and DNA from transcribing regions.

We find that during oogenesis, the transcribing loci from the fly genome are also covered by dP75, but likely via a distinct molecular mechanism. While still maintained the PWWP domain, dP75 lacks the AT hooks. However, it appears that its PWWP domain has binding affinity to both modified histones and DNA. When using H3K36me3 peptide as substrate, we detected weak but selective binding. Interestingly, although without the AT-hooks, our in vitro data indicate that the PWWP domain from dP75 can still recognize DNA, especially dT, revealing a new binding mode for this domain. It is worth noting that the 3’-untranslated regions from the *Drosophila* genome have a higher AT content, compared with coding regions (Zhang et al., 2004). And we indeed observed that dP75 has a higher affinity towards the 3’UTR regions. Thus, unlike its mammalian homolog which uses the PWWP and AT hooks coordinately to bind chromatin, the PWWP domain itself may facilitate the recruitment of dP75 to both modified histone and DNA simultaneously, resulting in the selection of targeting regions.

Via its C-terminus IBD-containing domain, dP75 recruits a putative histone kinase, Jil-1, to its binding sites. While whether Jil-1 indeed leads to histone phosphorylation on these specific loci is still to be determined, we found it functions together with dP75 to prevent the spreading of H3K9me2, thus likely maintaining local chromatin at an active state to ensure oogenesis. It appears that the dP75 and Jil-1 partnership is not only restricted within ovaries, previous mass spectrometry experiments also captured them in the same complex in S2 cells (Wang et al., 2013), which were likely derived from a macrophage-like lineage. Consistently, we also observed dP75 expression in somatic tissues, albeit at a slightly lower level compared with ovary. Interestingly, in somatic cells, Jil-1 also binds with transcribing regions, subsequently propelling transcription (Regnard et al., 2011). Given that the gene repertoire for active transcription is fundamentally different between germline and somatic tissues, the dP75-Jil-1 complex unlikely regulates the same set of targets in different cell types. This indicates that dP75 is recruited at the downstream of transcription initiation, rather than defining gene regions for activation. Thus, we propose that the general function of dP75-Jil-1 complex is protecting transcribing regions from the erosive spreading of H3K9me2, thus fine-tuning gene expression. This predicts that loss of dP75 or Jil-1 would not globally affect gene expression, but might lead to gene suppression when there is a high level of local H3K9me2. Our transcriptome data from dP75 or Jil-1 depleted ovaries indeed support this hypothesis.

Mice with compromised p75 expression suffer with fertility defects (Sutherland et al., 2006), but the underlying molecular mechanism is still unclear. Phylogenetic analysis shows that the closest mammalian homolog of Jil-1 is MSK1, which is also an H3S10 kinase and has been suggested to fine tune gene expression under stress conditions (Reyskens and Arthur, 2016). Interestingly, both p75 and MSK1 are highly expressed in mouse ovaries (unpublished data). Additionally, previous studies demonstrate that LEDGF/p75 binds with MLL1 complex (Mereau et al., 2013; Yokoyama and Cleary, 2008), and MLL1 is suggested to physically interact with MSK1 (Wiersma et al., 2016). Therefore, the partnership between dP75 and Jil-1 is likely to be conserved in mammals. Perhaps so does their molecular function on safeguarding oogenesis.

## MATERIALS AND METHODS

### REAGENTS

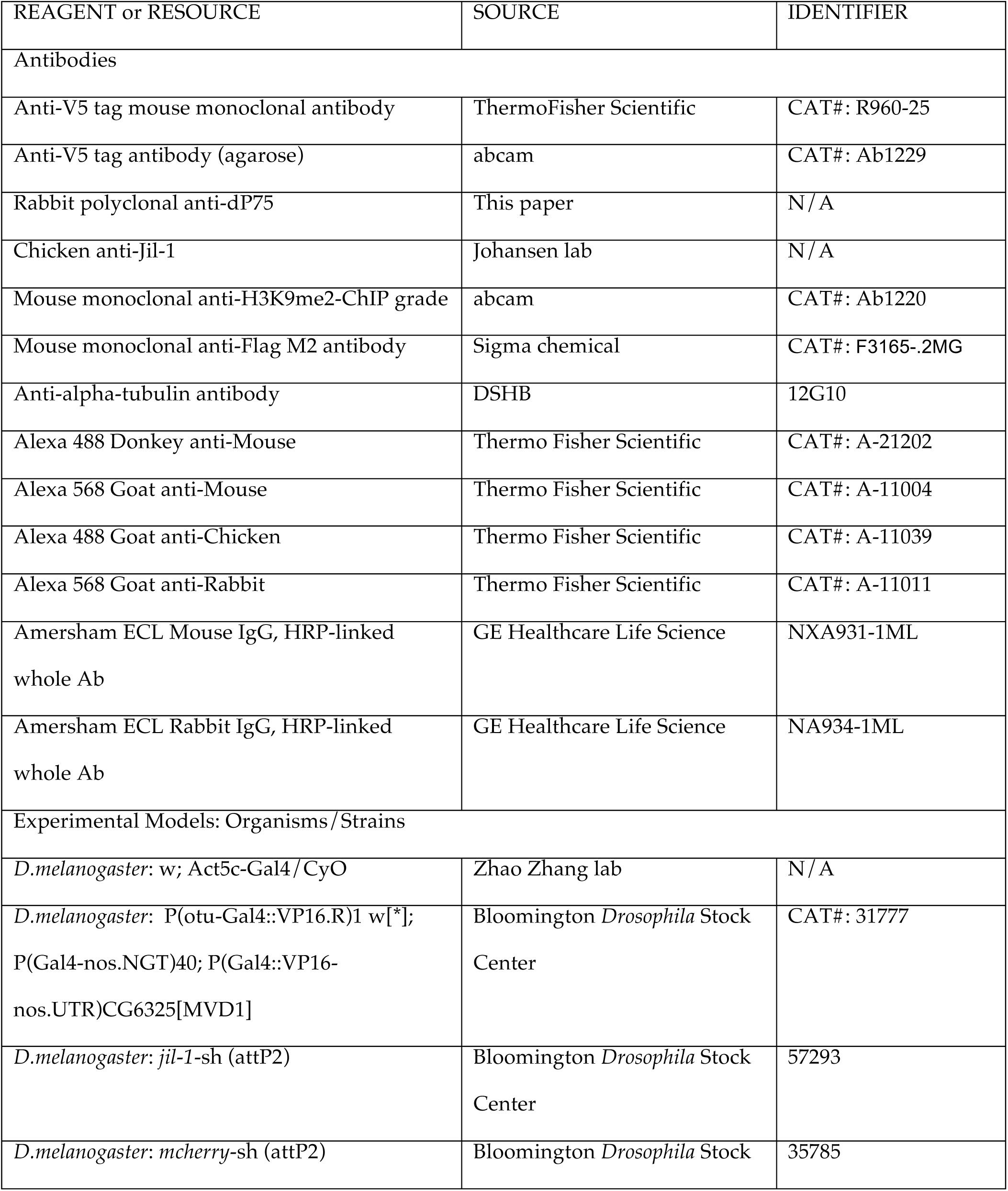
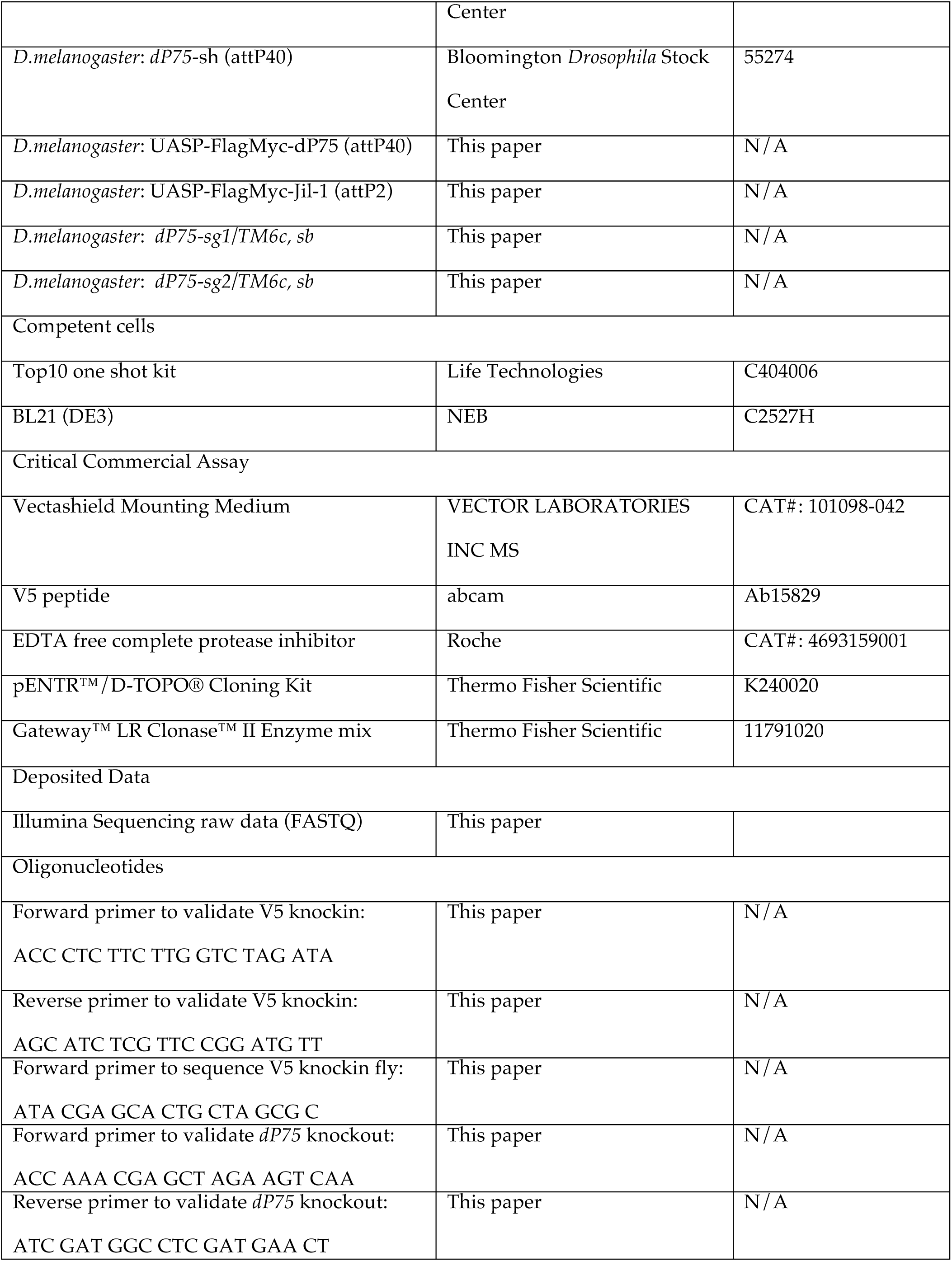
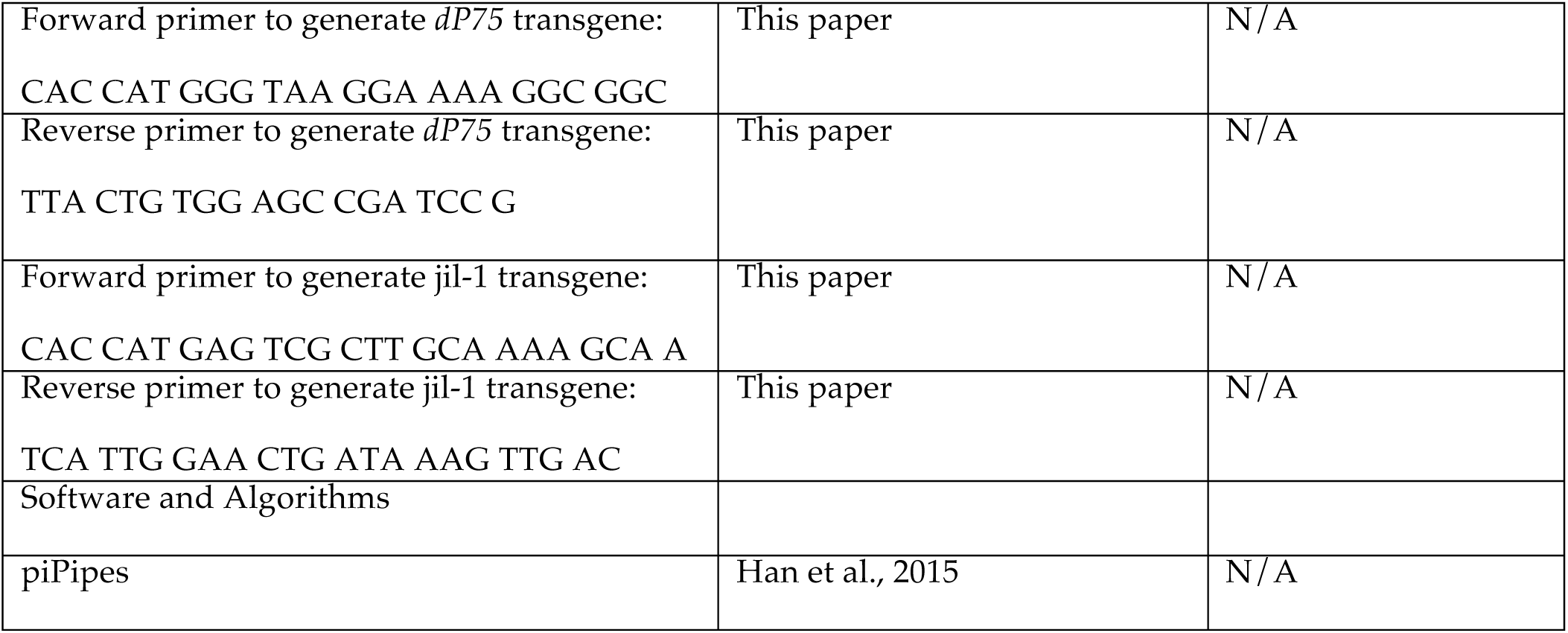

### Fly strains and husbandry conditions

The following fly lines were used: MTD-Gal4: P(otu-Gal4::VP16.R)1 w[*]; P(Gal4-nos.NGT)40; P(Gal4::VP16-nos.UTR)CG6325[MVD1], *mcherry*-sh (attP2) (Bloomington Stock Center), *jil-1*-sh (attP2) (Bloomington Stock Center), *dP75*-sh (attP40) (Bloomington Stock Center), *dP75^sg1^*/*dP75^sg2^* (this study), V5::dP75 (this study), UASp-Flag::Myc::Jil-1 (sttP2, this study), UASp-Flag::Myc::dP75 (attP40, this study). All fly strains were maintained at room temperature on standard agar-corn medium.

### Hatch ratio test

Four to six virgin females of 4-6-day-old with corresponding genotype, as well as equal number of wild type Origen R males were put together in an empty bottle, and molasses plate with a chunk of wet yeast paste on it was served each day to the flies. Molasses plates were collected after 24 hours to allow females to lay eggs on them, and maintained at room temperature for another 24-30 hours to allow eggs to hatch. Hatching rates were calculated by dividing the number of hatched eggs to number of total eggs laid. Four to six females were used for each genotype in each test, and at least two independent replicates were carried out for each experiment and reached the same conclusion.

### Generation of knockout and knockin flies with CRISPR/Cas9

For knockout flies, two separate sgRNAs were designed by using the webpage “http://tools.flycrispr.molbio.wisc.edu/targetFinder/index.php”. PCR sgDNAs by using pRB17 plasmid and sgRNA core as templates, and using U6_promo_F, sgRNA_U6term_R, and U6_yourguide_F together as primers. Note that sequence of the guide RNA I used was incorporated in the U6_yourguide_F primer. This overlapping PCR generated a single product of around 560 base pair, with U6 promoter followed by sgDNA sequence. The purified PCR products were sent out for Rainbow company for transgenic fly to inject the *y*,vas-Cas9 fly stain. PCR products of sgRNA for *white* were co-injected to the flies. The hatched flies from the injection was mated to balancer flies individually, and the second generation from each individual cross was first sorted based on eye color, followed by examination of whether our target gene *dP75* was disrupted using PCR and sequencing. Negative flies from the injection with the same background but without disruption of *dP75* were used as control animals.

To generate V5::dP75 flies, donor sequence was generated by annealing 800bp left arm homology sequnce, V5 tag sequence, and 600bp right arm homology sequence together by Gibson assembling, and cloning into pUC19 plasmid. Note that both PAM sequences corresponding to the two sgRNAs were mutated in the V5 knock-in construct to avoid cutting in the successful knock-in flies. Purified PCR products of two separate sgRNAs (as prepared above) and *white* sgRNA, as well as the donor plasmid were provided for injection of *y*,vas-Cas9 flies. Flies from the injection were sorted similar as described before for the knock-out injection, and followed by PCR and sequencing for the region of *dP75* to ensure knock-in fidelity. All the constructs, sequences, and successful knock-in and knock-out flies were identified by PCR and verified by sequencing corresponding regions.

### Immunostaining

Five pairs of ovaries from 4-8 day old flies were dissected in PBS, and immediately submerged in PBS with 4% formaldehyde for 12 minutes. Then ovaries were washed in PBST (PBS solution supplemented with 0.2% Triton X-100) for 3 times with a total of 30 minutes. Ovaries were blocked in PBST with 9% NGS overnight at 4 °C. Primary antibodies were added in blocking buffer after that, and ovaries were incubated for another 12 hours at 4 °C. Ovaries were washed for 3 times with a total of 30 minutes, and then incubated in blocking buffer with secondary antibody overnight at 4 °C. Ovaries were washed for another 3 times with a total of 30 minutes, followed by incubation in PBS with DAPI for 8 minutes. Ovaries were washed twice before mounting with 20 µl Vectashield Mounting Medium. Primary antibodies used in this study include: anti-dP75 (1:100), anti-V5 (1:100), anti-Jil-1 (1:100), anti-H3K9me2 (1:500).

### Affinity purification and mass spectrum analysis

Fifty pairs of fly ovaries from V5 knock-in flies at the endogenous *dP75* locus, as well as the same amount of wild type fly ovaries without V5 tag were dissected in cold PBS, and subjected to lysis with HEPES NP40 lysis buffer (50mM HEPES, KOH ph7.5, 150mM KCl, 3.2 mM MgCl2, 0.5% NP40, supplemented with Roche EDTA free complete protease inhibitor right before use). Sonicate the lysate in bioruptor on medium strength for 7 minutes (15 seconds on, 1 minute off). The supernatants were collected after spin at top speed, and subjected to immunoprecipitation. Forty µl V5 agarose beads were washed in lysis buffer and applied to each sample, followed by incubation of 3-5 hours at 4°C with gentle rotation. After IP, the beads are washed three times with the lysis buffer, and proteins were eluted with 40 µl lysis buffer supplemented with 150 µg/ml V5 peptide. The elutions were sent to Mass Spectrometry and Proteomics Facility of Johns Hopkins university for mass spectrum analysis.

### Antibody generation

Full-length dP75 protein with GST tag was purified from BL21 *E.coli* by using Glutathione Sepharose 4B GST-tagged protein purification resin (GE healthcare, 17075601) and following the protocol in GE healthcare webpage. The purified protein was sent to Pocono Rabbit Farm for polyclonal antibody production in rabbit. The final bleed showed highest efficiency and was used in western blot and immunostaining experiments.

### ChIP-Seq

Fifty pairs of fly ovaries were dissected in cold PBS for each sample, and fixed by paraformaldehyde at a final concentration of 1.8% for 10 minutes. Glycine were added to a final concentration of 125mM to stop fixation. The samples were washed with cold PBS and immersed in sonication buffer (50mM Tris-HCl pH8.0, 10mM EDTA, 1% SDS). Protease inhibitor was added right before use). Samples were sonicated with bioruptor, followed by spinning at top speed to remove the pellet. The lysates were diluted ten times, and 100 µl diluted lysate was saved as input, and the rest were used for IP. For IP, equal amount (25 µl) of washed protein A and protein G beads were first incubated with antibody for 3-4 hours at 4 °C. Then the lysates were applied to washed beads, and incubated at 4 °C overnight with gentle rotation. After IP, the beads were washed with low salt, high salt, LiCl buffer, and TE buffer sequentially. Direct elution buffer (10 mM Tris-HCl pH 8.0, 300 mM NaCl, 5 mM EDTA, and 0.5% SDS) was added, and samples were incubated at 65 °C for 6 hours for elution. The eluted DNA was extracted by phenol/chloroform, and was used for library preparation. The libraries were sequenced by Illumina NextSeq with single-end 75nt runs. The ChIP-Seq data were analyzed by piPipes (Han et al., 2015) and deep Tools, and the same number of mapped reads was used for comparison between control and experimental groups.

### mRNA sequencing

Total RNAs were extracted from six pairs of 4-6 day old fly ovaries of each sample, followed by ribosomal RNA depletion with RiboZero magnetic kit and Turbo DNase treatment. These RNAs were purified by RNA Clean & Concentrator-5 kit. The purified RNAs were subjected to library preparation by following the protocol described (Zhang et al., 2012). Libraries were sequenced with Illumina NextSeq with paired-end 75nt runs. The RNA-Seq data were analyzed by piPipes (Han et al., 2015) and hisat2. During plotting process, few genes with fpkm larger than 13 are not shown in the plot since none of them showed altered expression by loss of either Jil-1 or dP75.

### Plasmid construction

For UASP-Flag::Myc::dP75 construct, full length dP75 was amplified from fly cDNA, and cloned to gateway entry vector, followed by recombination to UASP-Flag:Myc gateway destination vector with the attB sites. The positive colonies were validated by sequencing. The UASP-Flag::Myc::Jil-1 was constructed in similar way. For in vitro assays, DNA fragment encoding the PWWP domain of dP75 (residues 8-91) was cloned into the pET28-SMT3 vector between the *BamH*I and *Sal*I sites, which contains an N-terminal Ulp1 cleavable His6-SUMO tag. *dP75* plasmid with mutations were generated by Site-Directed Mutagenesis Kit (NEB). For yeast two-hybrid assays, fragments of dP75 (residues 1-475, 1-90, 85-300, 211-475, 209-445, 423-475) were cloned between *BamH*I and *Sal*I sites of the pBTM116 vector (Clontech). Fragments of Jil-1 (residues 1-1207, 1-255, 255-541, 525-990, 984-1122, and 1100-1207) were cloned between *BgII*I and *Xho*I sites of the pACT2 vector (Clontech). All the mutants were generated using a Site-Directed Mutagenesis Kit (NEB) according to the manufacturer’s instructions. All constructs were sequence verified.

### Protein expression and purification

For in vitro assays, His_6_-SUMO-tagged recombinant proteins were expressed in *Escherichia coli* BL21 (DE3) under kanamycin selection. Cells were grown to an OD_600_ of 0.6, and protein expression was induced with 0.2 mM IPTG overnight at 18°C. The cells were collected by centrifugation at 4,000 rpm for 15 minutes and were lysed by French Press (JNBIO) at 4 °C. His6-SUMO-tagged proteins were purified by affinity chromatography using a HisTrap HP column (GE Healthcare), followed by the cleavage of the His_6_-SUMO tag using Ulp1 protease. The proteins were further purified by ion-exchange chromatography using a HiTrap S column (GE Healthcare) and eluted in a gradient of NaCl. Size-exclusion chromatography was performed for the final purification of wild-type and mutant dP75 PWWP using Superdex G75 Hiload 16/60 column (GE Healthcare). Fractions containing target proteins were pooled and concentrated to 10 mg/ml in a buffer containing 10 mM Tris-HCl, pH 8.0, 100 mM NaCl, and 1 mM DTT.

### Isothermal titration calorimetry (ITC)

DNA fragments used in ITC were synthesized (Sangon). dP75 PWWP as well as DNA fragments or histone peptides were dialyzed overnight in a buffer containing 10 mM Tris, pH 8.0, 100 mM NaCl. ITC measurements were performed in duplicates at 20 °C, using an iTC200 (MicroCal, Inc). The samples were centrifuged before the experiments to remove any precipitates. Experiments were carried out by 20 injections of 2 μL of 2 mM DNA solution into the sample cell containing 0.2 M of dP75 PWWP in the dialysis buffer. Experiments were carried out by 20 injections of 2 μL of 1 mM wild-type or mutant Jil-1 peptides (1102-1112, 984-1004, 984-1004 (WT, F989A, G991V, F992A)) into the sample cell containing 0.1 mM of dP75 IBD-containing C-terminus region (residues 209-445) in the dialysis buffer. Binding isotherms and standard deviation were derived from nonlinear fitting using Origin 8.0 (MicroCal, Inc). The initial data point was routinely deleted. The ITC data were fit to a one-site binding model in 1:1 binding mode.

### Yeast two-hybrid assay

The yeast two-hybrid assays were performed with L40 strains containing pBTM116 and pACT2 fusion plasmids. The colonies containing both plasmids were selected on Leu-Trp-plates. To determine the strength of protein–protein interactions, β-galactosidase solution assays were performed using ortho-nitrophenyl-β-galactopyranoside (ONPG) as the substrate. The cells were inoculated into Leu-Trp-media and grown at 30 °C with vigorous shaking until the OD_600_ reached 0.6-1. At least three independent clones were selected. The cells (2mL) were harvested, centrifuged, and resuspended in 0.1 mL Z buffer (16.1 g/L Na_2_HPO_4_·7H_2_O, 5.50 g/L NaH_2_PO_4_·H_2_O, 0.75 g/L KCl, 0.246 g/L MgSO_4_·H_2_O, pH 7.0), frozen in liquid nitrogen, and thawed at 37 °C in a water bath. Repeat this step for two more times to lysis cells completely. Then the reaction systems (0.2 mL 4mg/mL ONPG in Z buffer and 0.9 mL 0.27% β-mercaptoethanol in Z buffer per 0.1 mL yeast cell resuspensions) were placed in a 30 °C incubator. After the yellow color developed, 0.5 mL 1 M Na_2_CO_3_ was added to the reaction and blank tubes. Reaction time was recorded in minutes. Reaction tubes were centrifuged at 14,000 g for 10 minutes and supernatants were carefully transferred to clean cuvettes and OD_420_ of the samples relative to the blank was recorded. At last, the β-galactosidase units were calculated using the following formula: β-galactosidase units = 1000×OD_420_/(t×V×OD_600_), where t is the reaction time (in minute) of incubation, V is the volume and in this case V=2, OD_600_ is the cell density of the culture.

### RNA FISH

The RNA FISH experiments were carried out according to the protocol described in *Wang et al., 2018*, *Cell*. Briefly, Stellaris RNA FISH probes were designed and purchased from LGC Bioresearch Technology. 3-5 pairs of ovaries were dissected in cold PBS, and fixed in 4% formaldehyde for 20 minutes. Ovaries were washed once with PBST and twice with PBS, followed by immersing in 70% (v/v) ethanol for 8 hrs at 4 °C. Then ovaries were washed with Wash Buffer A (LGC Biosearch Tech, Cat# SMF-WA1-60) at room temperature for 5 minutes, and incubated with 50 µl Hybridization Buffer (LGC Biosearch Tech, Cat# SMF-HB1-10) containing RNA probe set overnight at 37 °C. After that, ovaries were washed twice with Wash Buffer A at 37 °C for 30 minutes and once with Wash Buffer B (LGC Biosearch Tech, Cat# SMF-WB1-20) at room temperature for 5 minutes. Samples were mounted after final wash.

## QUANTIFICATION AND STATISTICAL ANALYSIS

The number of animals examined is indicated in materials and methods. Student’s T test was carried out, and the results are shown as mean ± standard deviation (SD). The biostatistics of deep-sequencing libraries is listed in Table S2.

## DATA AND SOFTWARE AVAILABILITY

Deep sequencing data are deposited to NCBI.

## ACKNOWLEDGMENTS

We thank the Johansen lab for providing Jil-1 antibody. We thank members from ZZ lab, Spradling lab, and Bortvin lab for valuable suggestions. We thank Dr. Wanbao Niu for sharing unplublished RNA sequencing data from mouse ovary. We thank Dr. Allan Spradling and Dr. Robert Levis for proof-reading of the manuscript. We thank Shanghai Science Research Center and Zhangjiang Laboratory for their instrumental support and technical assistance. This work was supported by the grant from the NIH to Z.Z. (DP5OD021355), as well as the National Natural Science Foundation of China (91640102 and 31870741 to Y.H.).

## AUTHOR CONTRIBUTIONS

K.D and Z.Z. conceived the project. K.D., Z.Z. and Y.H. designed the experiments. K.D. performed experiments regarding to flies. Y.L. and Y.Z. carried out the *in vitro* interaction assays and yeast two hybrid experiments. C.W. designed and constructed the mosaic analysis assay. K.D. analyzed the sequencing data. K.D., Y.H., and Z.Z. wrote the manuscript.

---

**Figure S1.**
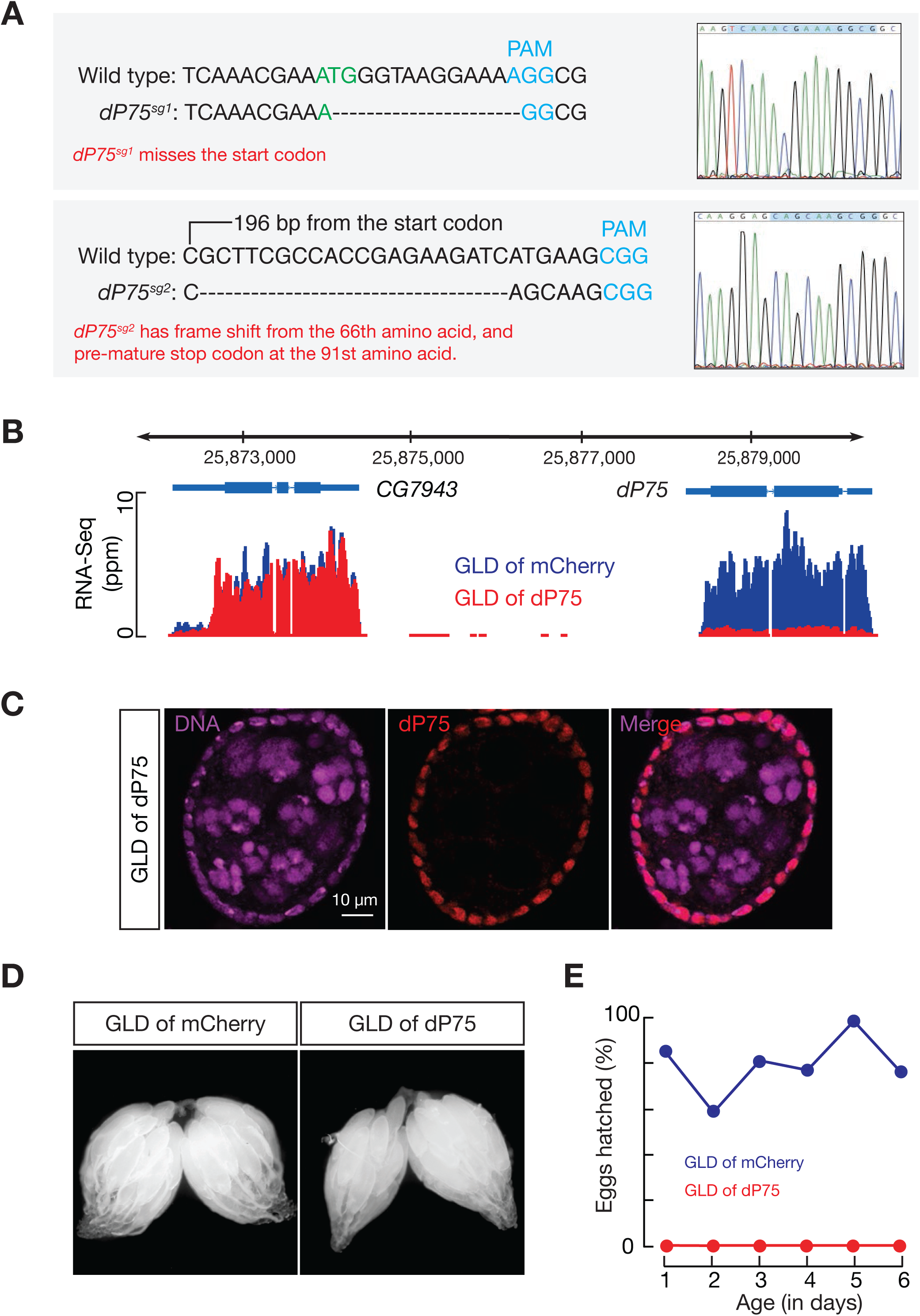
dP75 depletion in germ cells leads to female sterility, Related to Figure 1. (A) dP75 mutation alleles generated by two sgRNAs. Sequencing results from the homozygous fly alleles are shown on the right. (B) RNA-Seq signals for dP75 in fly ovaries upon depleting dP75 in germ cells. (C) Immunostaining of dP75 to test the efficiency of dP75 depletion in germ cells. (D) Ovary morphology appears as normal upon dP75 depletion in germ cells. (E) Depleting dP75 in germ cells leads to animal sterility. GLD = Germline Depletion.

**Figure S2.**
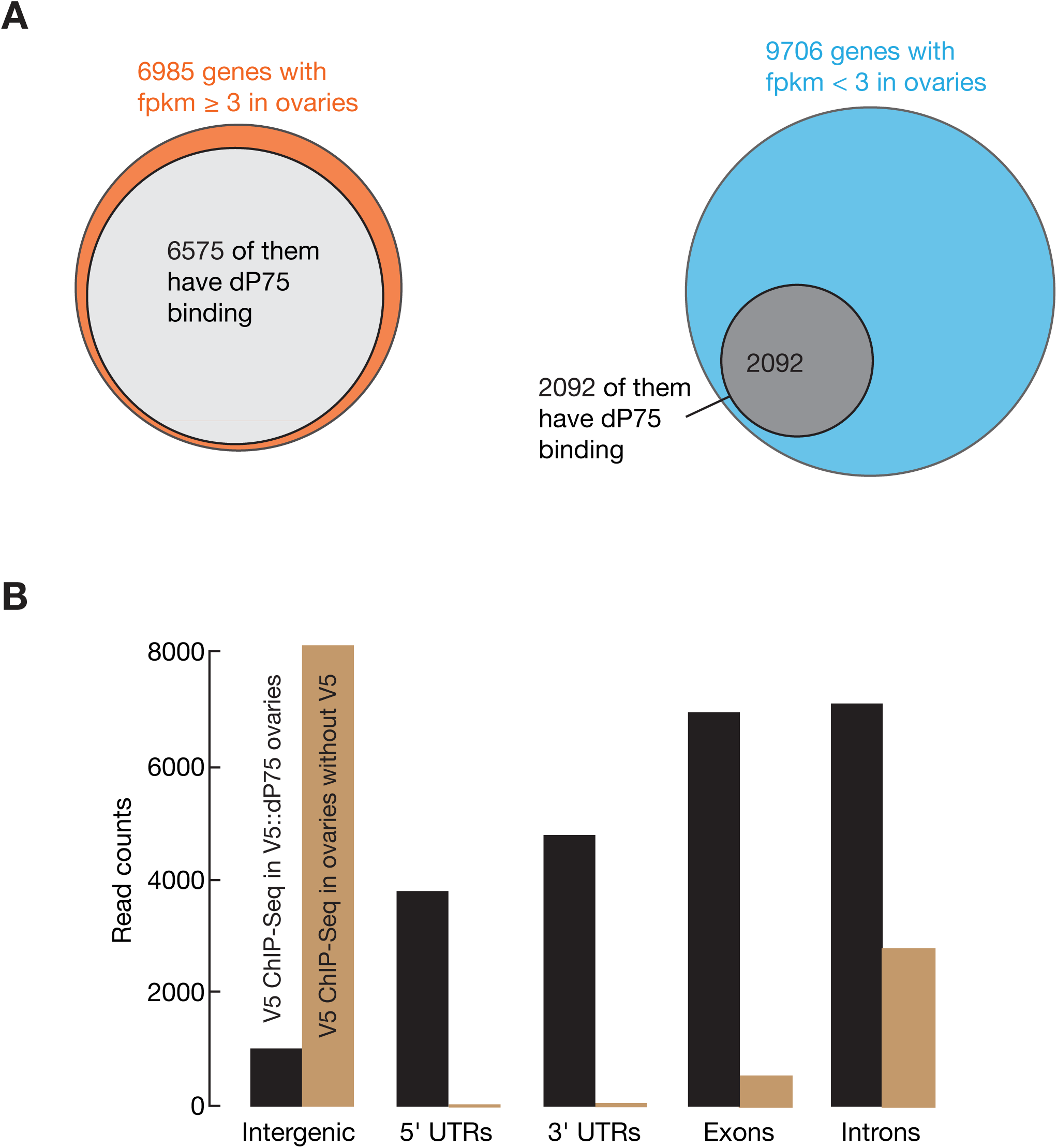
dP75 binds to chromatin regions with active transcription. (A) Venn diagrams showing the numbers of genes that are coated by dP75 having fpkm >= 3 (left panel) or having fpkm < 3 (right panel). (B) Bar graph showing the ChIP-Seq enrichment of dP75 at different genic regions.

**Figure S3.**
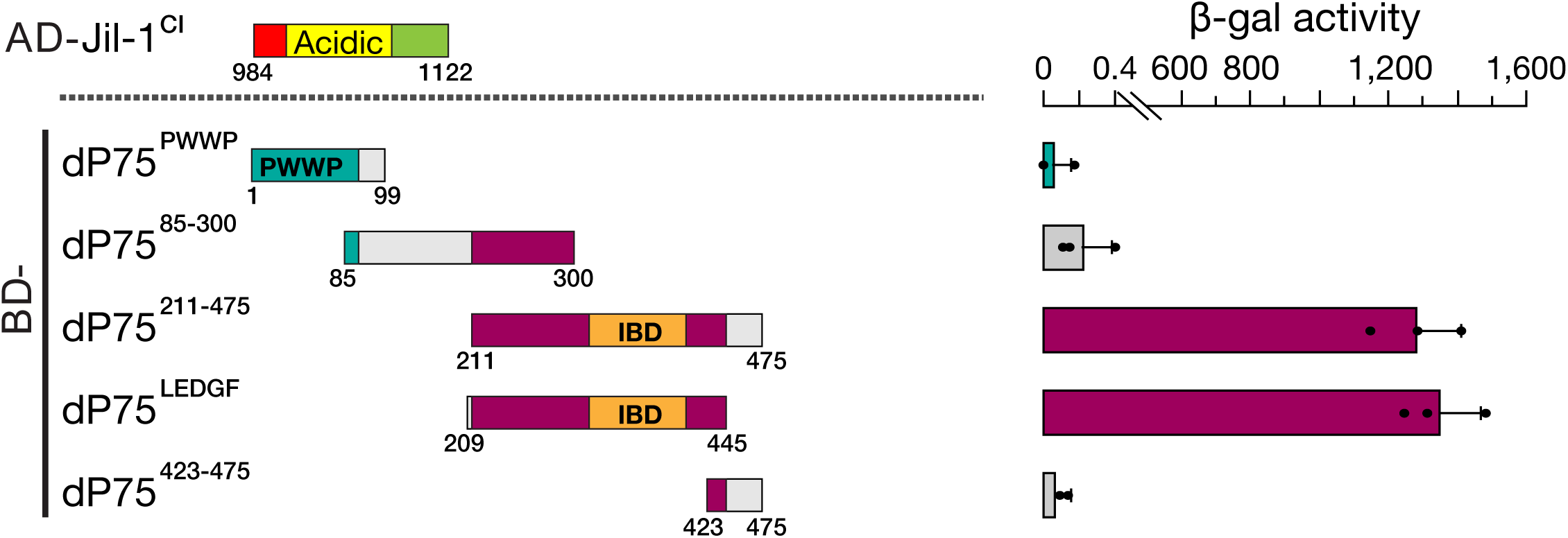
dP75 interacts with Jil-1 in Yeast-2-Hybrid assay. Interaction measurement between Jil-1C1 (C-terminus fraction 1 of Jil-1) and different truncated versions of dP75. The C-terminus fragment of dP75 containing the IBD domain binds with Jil-1C1 with the highest affinity.

**Figure S4.**
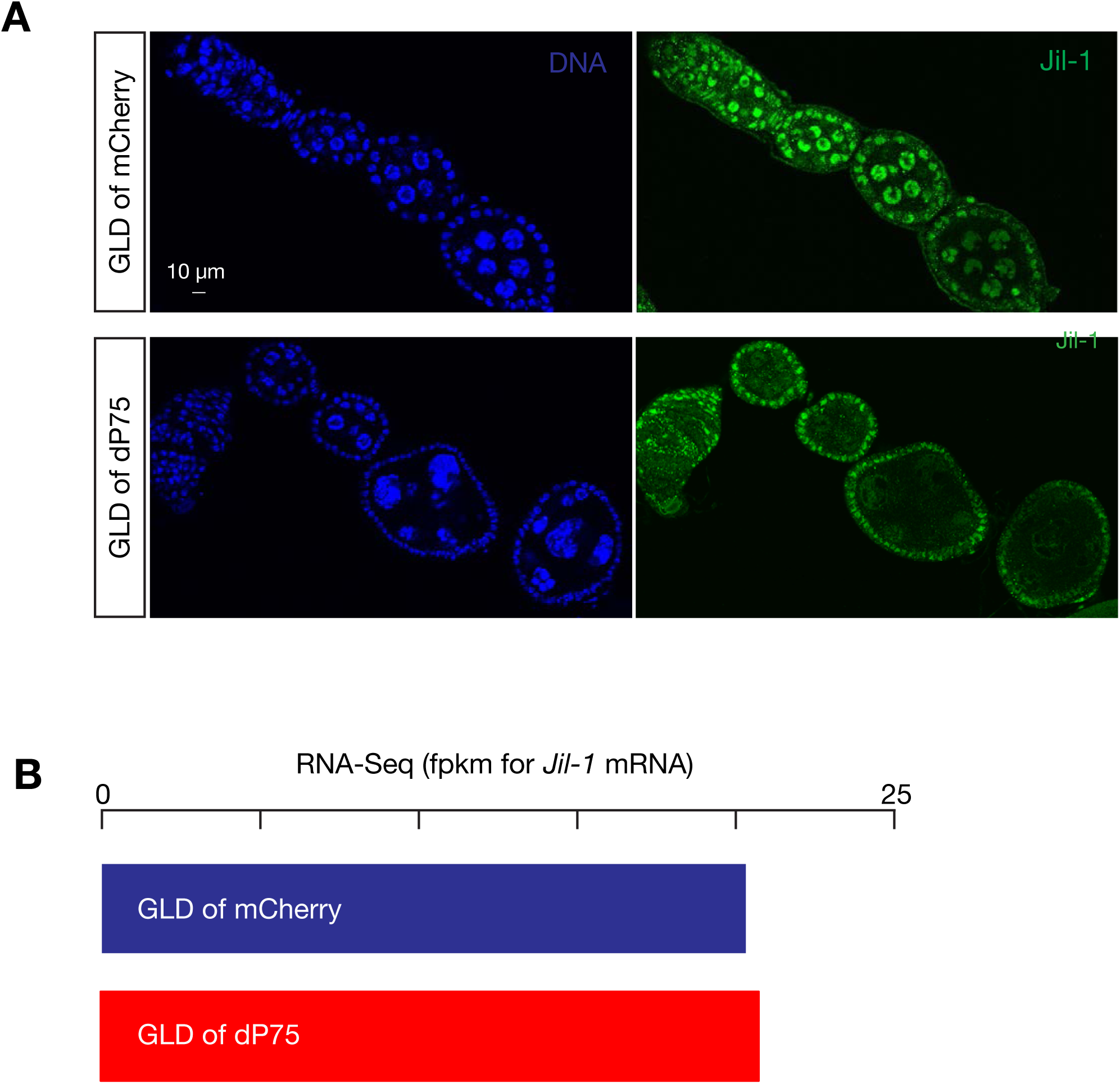
dP75 stabilizes Jil-1 in vivo. (A) Immunostaining of Jil-1 in control ovaries or ovaries with dP75 depletion from germ cells. (B) mRNA level of jil-1 is unaltered by dP75 depletion. GLD = Germline Depletion.

**Figure S5.**
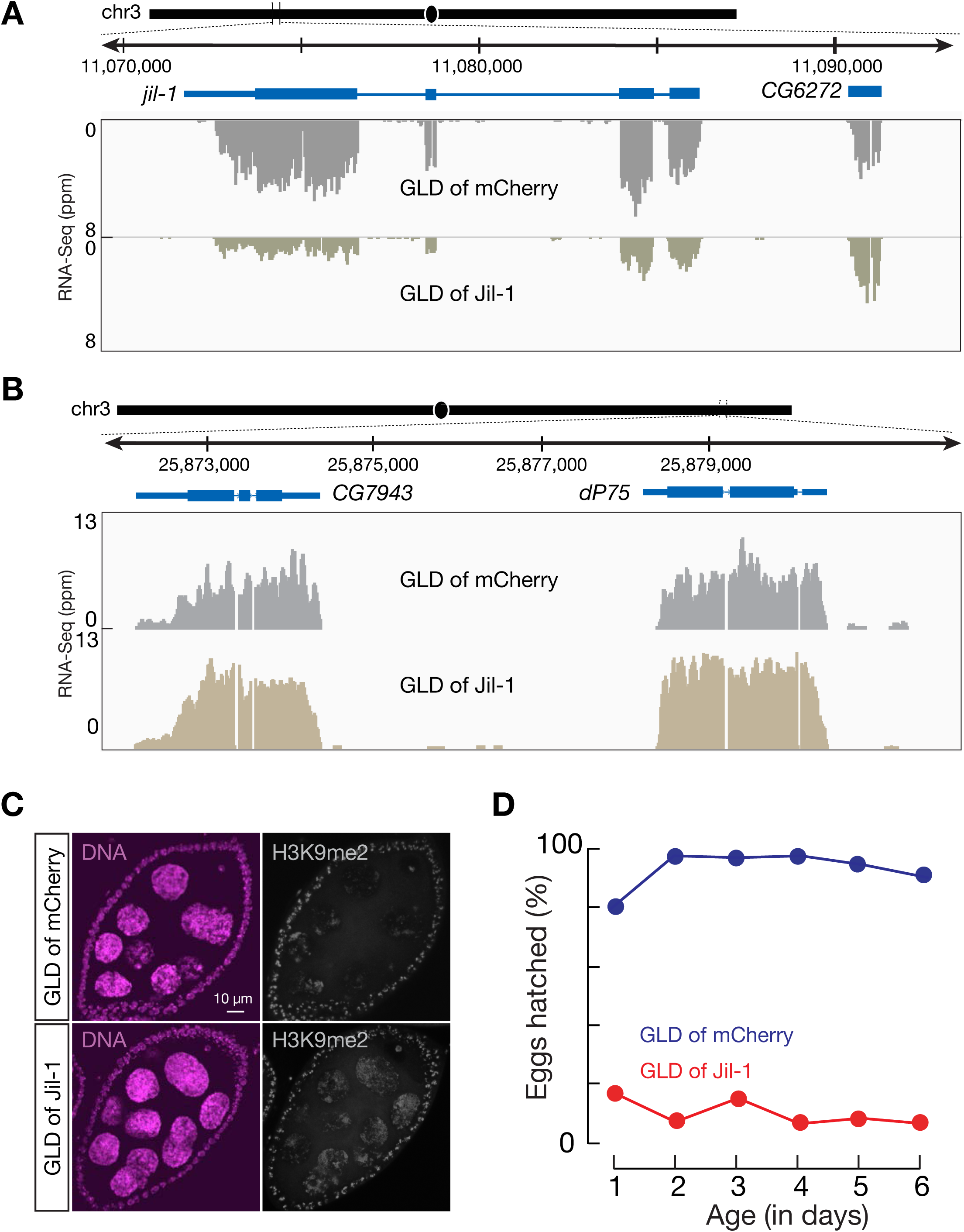
Germline depletion of Jil-1 results in H3K9me2 spreading and female sterility. (A) Efficiency of Jil-1 depletion in germ cells. Jil-1 mRNAs from somatic follicle cells are unaltered, likely contributing to the signal from the browser. (B) The mRNA level of dP75 is not affected by germline depletion of Jil-1. (C) Immunostaining of H3K9me2 in control ovaries or in ovaries with germline depletion of Jil-1. (D) Hatch ratio of control females and females with Jil-1 depletion in germ cells. GLD = Germline Depletion.

**Figure S6.**
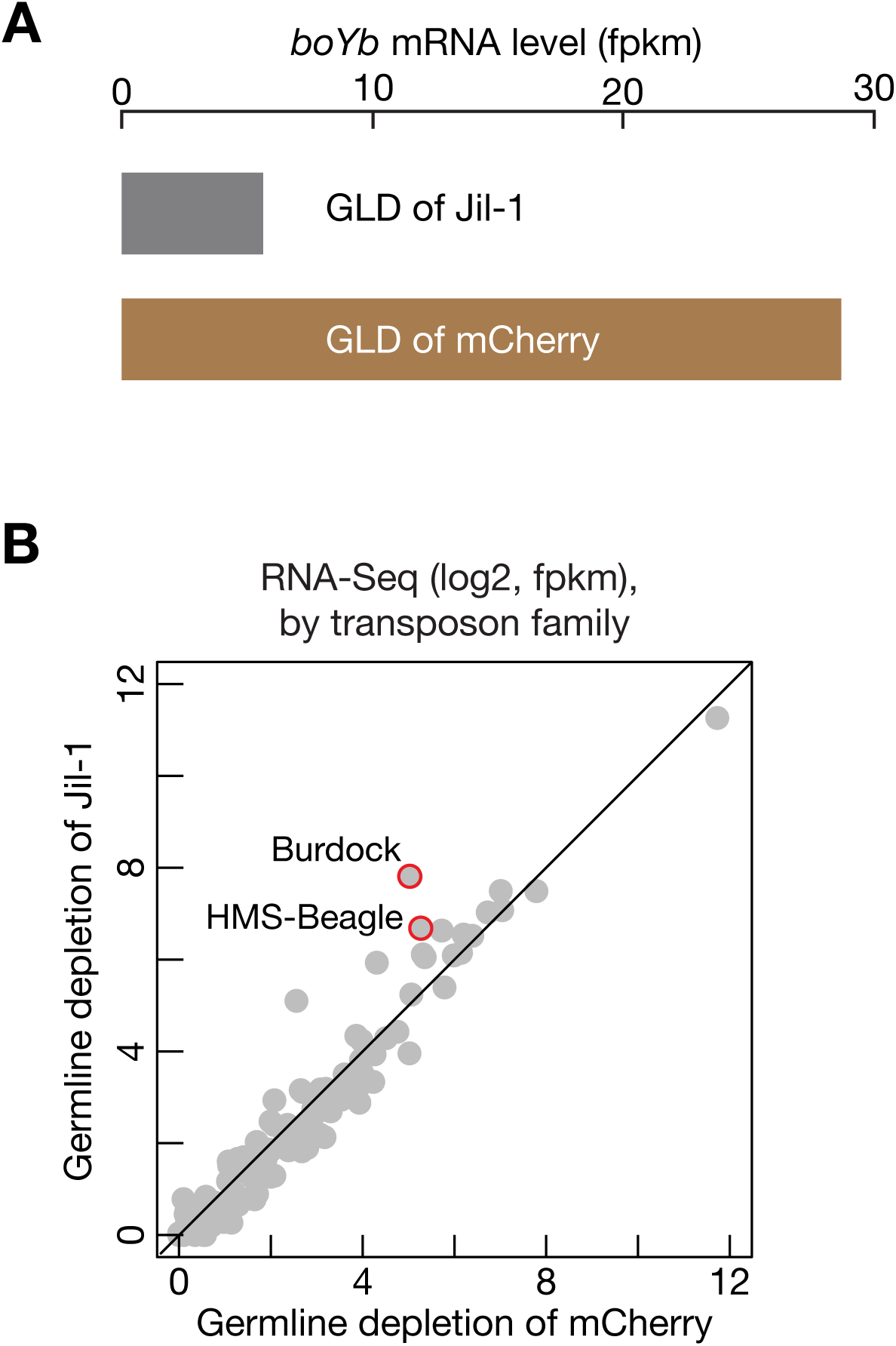
dP75 and Jil-1 are required for transposon suppression. (A) Depleting Jil-1 in germ cells leads to lower boYb expression. (B) The change in transposon expression, relative to control, was compared for ovaries with Jil-1 depleted in germ cells.

**Table S1.** Summarization of oogenesis defects in ovaries upon loss of dP75 in germ cells.

|  | **GLD of mCherry** | **GLD of dP75** |
| --- | --- | --- |
| **Abnormal sized egg chamber** | 6%<br>(2/33 ovarioles) | 50%<br>(30/60 ovarioles) |
| **More than 15 nurse cells** | 0%<br>(0/40 st6-8 egg chambers) | 5%<br>(2/40 st6-8 egg chambers) |
| **Blob-like chromosomes in st7-9** | 0%<br>(0/41 st7-9 egg chambers) | 71%<br>(47/66 st7-9 egg chambers) |
| **Smaller nurse cells in st7-9** | 6%<br>(7/112 st7-9 egg chambers) | 88%<br>(90/102 st7-9 egg chambers) |
| **Stretched oocyte DNA in st7-8** | 2%<br>(1/56 st7-8 egg chambers) | 27%<br>(15/55 st7-8 chambers) |

**Table S2.** Deep-sequencing biostat.

| Number | Sample name | Fragments number | Seq type | Mappability to fly genome |
| --- | --- | --- | --- | --- |
| 1 | GLD dP75 replicate 1 | 60370015 | RNA-seq | 89.64% |
| 2 | GLD dP75 replicate 2 | 127140079 | RNA-seq | 87.11% |
| 3 | GLD mCherry replicate 1 | 7744016 | RNA-seq | 87.56% |
| 4 | GLD mCherry replicate 2 | 154908012 | RNA-seq | 90.67% |
| 5 | GLD Jil-1 replicate 1 | 64000394 | RNA-seq | 79.16% |
| 6 | GLD Jil-1 replicate 2 | 59348823 | RNA-seq | 77.27% |
| 7 | GLD mCherry control replicate 1 | 75387201 | RNA-seq | 74.45% |
| 8 | GLD mCherry control replicate 2 | 54199871 | RNA-seq | 72.59% |
| 9 | wild-type replicate 1 | 56624560 | RNA-seq | 93.16% |
| 10 | wild-type replicate 2 | 50092614 | RNA-seq | 88.41% |
| 11 | dP75 mutant replicate 1 | 61293162 | RNA-seq | 93.90% |
| 12 | dP75 mutant replicate 2 | 60944788 | RNA-seq | 88.48% |
| 13 | No V5 tag ChIP-seq | 47909827 | ChIP-seq | 77.63% |
| 14 | No V5 tag ChIP-seq input | 78708466 | ChIP-seq | 89.60% |
| 15 | V5-dP75 ChIP-seq | 46504024 | ChIP-seq | 91.29% |
| 16 | V5-dP75 ChIP-seq input | 85287523 | ChIP-seq | 89.39% |
| 17 | wild-type H3K9me2 ChIP-seq replicate 1 | 55682788 | ChIP-seq | 72.73% |
| 18 | wild-type H3K9me2 ChIP-seq input replicate 1 | 74708824 | ChIP-seq | 87.56% |
| 19 | wild-type H3K9me2 ChIP-seq replicate 2 | 74821158 | ChIP-seq | 76.84% |
| 20 | wild-type H3K9me2 ChIP-seq input replicate 2 | 89226713 | ChIP-seq | 86.10% |
| 21 | dP75 mutant H3K9me2 ChIP-seq replicate 1 | 74659995 | ChIP-seq | 85.39% |
| 22 | dP75 mutant H3K9me2 ChIP-seq input replicate 1 | 75551897 | ChIP-seq | 89.73% |
| 23 | dP75 mutant H3K9me2 ChIP-seq replicate 2 | 72270441 | ChIP-seq | 81.15% |
| 24 | dP75 mutant H3K9me2 ChIP-seq input replicate 2 | 45292235 | ChIP-seq | 86.01% |

Note: 1. Gene expression from RNA-seq was calculated based on FPKM (Fragments per kilobase per million reads). 2. The same number of mapped reads for ChIP-seq were used when comparing experimental group and control group.

